# Hierarchical Classification of Insects with Multitask Learning and Anomaly Detection

**DOI:** 10.1101/2023.06.29.546989

**Authors:** Kim Bjerge, Quentin Geissmann, Jamie Alison, Hjalte M. R. Mann, Toke T. Høye, Mads Dyrmann, Henrik Karstoft

**Affiliations:** Department of Electrical and Computer Engineering, Aarhus University,Finlandsgade 22, 8200 Aarhus N, Denmark; Center for Quantitative Genetics and Genomics, Aarhus University, C.F. Møllers Allé 3, 8000 Aarhus C, Denmark; Department of Ecoscience and Arctic Research Centre, Aarhus University, C.F. Møllers Allé 4-8, 8000 Aarhus C, Denmark

**Author notes:** Corresponding author *Email address:* (Kim Bjerge).

**Keywords:** `A`nomaly detection, Computer Vision, Deep Learning, Hierarchical classification, Insects, Taxonomy

## Abstract

Cameras and computer vision are revolutionising the study of insects, creating new research opportunities within agriculture, epidemiology, evolution, ecology and monitoring of biodiversity. However, a major challenge is the diversity of insects and close resemblances of many species combined with computer vision are often not sufficient to classify large numbers of insect species, which sometimes cannot be identified at the species level. Here, we present an algorithm to hierarchically classify insects from images, leveraging a simple taxonomy to (1) classify specimens across multiple taxonomic ranks simultaneously, and (2) highlight the lowest rank at which a reliable classification can be reached. Specifically, we propose multitask learning, a loss function incorporating class dependency at each taxonomic rank, and anomaly detection based on outlier analysis for quantification of uncertainty. First, we compile a dataset of 41,731 images of insects, combining images from time-lapse monitoring of floral scenes with images from the Global Biodiversity Information Facility (GBIF). Second, we adapt state-of-the-art convolutional neural networks, ResNet and EfficientNet, for the hierarchical classification of insects belonging to three orders, five families and nine species. Third, we assess model generalization for 11 species unseen by the trained models. Here, anomaly detection is used to predict the higher rank of the species not present in the training set. We found that incorporating a simple taxonomy into our model increased accuracy at higher taxonomic ranks. As expected, our algorithm correctly classified new insect species at higher taxonomic ranks, while classification was uncertain at lower taxonomic ranks. Anomaly detection can effectively flag novel taxa that are visually distinct from species in the training data. However, five novel taxa were consistently mistaken for visually similar species in the training data. Above all, we have demonstrated a practical approach to hierarchical classification based on species taxonomy and uncertainty during automated *in situ* monitoring of live insects. Our method is simple and versatile and could be implemented to classify a wide range of insects as well as other organisms.

## 1. Introduction

Insects are the most diverse group of animals and live in almost every habitat. They play pivotal roles in most ecosystems and have considerable agronomic importance (e.g., as pests and pollinators). Technologies such as automated monitoring with computer vision and deep learning are revolutionizing insect surveys (van Klink et al., 2022; Lima et al., 2020). In particular, computer vision methods have shown great promise, both in real time (Bjerge et al., 2021b,a) and offline, on images acquired by time-lapse (TL) cameras (Geissmann et al., 2022; Bjerge et al., 2023). However, images of insects using fixed position cameras are frequently not offering sufficient details for species level identification. Typically, the insect will make up a small part of a high resolution image. As such, it will be important to incorporate a hierarchy in the taxonomic assignment of insect identification from images to accommodate for this uncertainty.

Automation is essential for spatiotemporal upscaling of insect monitoring globally and for efficient pest management (Preti et al., 2021). Deep-learning methods are increasingly used for classification in many fields and are emerging in entomology (Høye et al., 2021). Such methods require large training datasets for prediction robustness, which has so far been a limiting factor in entomology. Specifically, insects are very diverse and their abundance is highly variable, which often results in unbalanced datasets with numerous classes nested inside each other in a taxonomic hierarchy. As these methods mature and are implemented in monitoring programs, the demand for greater taxonomic resolution of the classifications will increase. So far, the level of fine-grained classification is still rather limited, not least because classification models rely on training data where images are assigned to species labels.

### 1.1. Background and related work

This section outlines different approaches to classification of insects with deep learning, including hierarchical classification, applied in different domains. Multitask learning adapted for hierarchical classification is introduced to model the taxonomic hierarchy within the network architecture. Finally, anomaly detection is discussed to identify uncertain or previously unseen species not included in the training of the hierarchical classification model.

Convolutional neural networks (CNNs) are a widely used deep learning tool to detect and classify insects and other organisms. It is most common to use a traditional “flat” classification, whereby deep learning models do not explicitly consider evolutionary relationships between taxa (Hansen et al., 2020; Kasinathan et al., 2021; Kittichai et al., 2021; Xia et al., 2018). However, those relationships are hierarchical in nature, and hierarchical classification has been researched in different application domains (Silla and Freitas, 2011; Salakhutdinov et al., 2013; Park and Kim, 2020) such as diatom images (Dimitrovski et al., 2012), disease detection (An et al., 2021) and protein families (Sandaruwan and Wannige, 2021). For animals such as arthropods (Tresson et al., 2021) and fish species (Gupta et al., 2022) hierarchical classification has been investigated with the object detector YOLOv3 (Redmon and Farhadi, 2018) designed to detect and classify species using a “flat” multi-class structure. Tresson et al. (2021) used a two-step approach with a YOLOv3 object detector to detect and classify arthropods in five super-classes, using a separate model to classify cropped images at the species level. Similarly, and with application to the classification of insects, Ong and Hamid (2022) classified museum insect specimens into their respective taxonomic ranks (i.e., order, family and genus) by training a different CNN for each rank. The above studies enable classification of specimens at multiple taxonomic ranks. However, training a separate model for each taxonomic rank means that hierarchical information is not shared between models, and is not fully leveraged.

Hierarchical classification can be applied to a traditional “flat” multi-label network within a single model, and may improve predictive performance at multiple levels of the hierarchy. This is done by using a class label for each level of the hierarchy. To handle the hierarchy dependency, a loss function is introduced such that a higher penalty is assigned for wrong predictions in the multi-label hierarchy. Wu et al. (2019) propose a hierarchical cross-entropy (HXE) loss function to organize a “flat” network of leaves in a tree organized into a semantically meaningful hierarchy. However, they assess their method on text classification, where performance varies across datasets and the hierarchical loss function is only for “flat” networks which is not an optimal way to model a hierarchy. Bertinetto et al. (2020) propose modifications of the cross-entropy loss to handle class hierarchy dependency in “flat” networks. Both methods only focus on the loss function and do not incorporate the class hierarchy into the CNN architecture.

Hierarchical classification in a single model where the CNN architecture reflects the hierarchy with a dependency loss function is proposed by (Gao, 2020; La Grassa et al., 2021). Gao (2020) present a method based on deep hierarchical classification for category prediction in E-commerce systems that uses a CNN as its backbone with embedding branches to handle each level in the categorical hierarchy. The method is noteworthy since it takes advantage of state-of-the-art CNN architecture but replaces the output layers with hierarchical embedding branches. Further, the authors define a novel combined loss function to punish hierarchical prediction losses. La Grassa et al. (2021) propose an architecture that can extend any CNN model with a stack of linear layers using cross-entropy combined with a centre-loss function. However, experiments show that the centre-loss function will perform poorly together with anomaly detection since the variance of output distribution will be small and it will thereby be difficult to identify outliers.

Multitask learning (MTL) is the process of training a model to learn several related tasks, which is relevant in the context of hierarchical classification, and has never been used to solve classification challenges for biological organisms. MTL improves generalization by inductive information transfer and by using the domain information contained in the training data of related tasks. It does this by learning tasks in parallel while using a shared representation; what is learned for each task can help other tasks (Maurer et al., 2016).

Hard parameter sharing is the most commonly used approach by which the hidden CNN layers are shared between all tasks while keeping several task-specific outputs (Zhang et al., 2022). Caruana (1997) present examples of the usages of MTL. One example was to predict steering directions for road-following vehicles. Based on road images, the model would predict, for example, the number of road lanes, the position on the road, the intensity of the road surface and regions boarding the road. Taylor et al. (2020) used MTL on clinical data to predict, for example, mood, stress, and health of people across different populations. The new approach in our work is to apply MTL for hierarchical classification in a single model by adding a dependency loss function between layers.

Identifying cases of uncertainty or novelty is vital in many deep learning applications, particularly in hierarchical classification. Anomaly detection, also known as outlier detection or novelty detection (Pang et al., 2021), can help to achieve this by finding patterns in data that do not adhere to the expected norms, given previous observations. Most deep semi-supervised approaches first model normal behaviour to later identify anomalies (Villa-Pérez et al., 2021). The distribution of normal observations is learned based on the output scores assigned by the trained CNN model. A threshold rule is then applied to tag anomalies for scores below a defined threshold value outside the learned distribution. Since it is unfeasible to collect data to train models that can classify all insect species, we suggest using anomaly detection embedded in the classification model to identify uncertain or previously unseen species. This concept, used in our work, is called threshold-based anomaly tagging.

### 1.2. Contribution

In this work, we propose hierarchical classification as an approach to classify rare and unseen species at higher taxonomic ranks (e.g. genus, family or order). We propose a method to hierarchically classify insect species according to their biological taxonomy inspired by Gao (2020) and La Grassa et al. (2021). Instead of embedding branches or a stack of linear layers, we use the architecture from multitask learning with parallel, fully connected layers for each level in the hierarchy.

Insects (*Insecta*) are a ‘class’ of organisms, where class is a rank in the traditional Linnaean taxonomy. Our method uses multitask and transfer learning to classify insects at three different taxonomic ranks: order, family, and species. This work aims to robustly detect previously unseen species, yet classify them to their correct higher taxonomic rank. Such species should be detected as ‘unsure’ (anomalies), at the species level, but classified correctly at higher taxonomic rank. We detect anomalies as outliers from learning the distribution of the output for each rank in the hierarchy of the trained model.

We hypothesize that it is possible to classify insect species in taxonomic ranks and detect unsure insect taxa with the trained model as anomalies. We assess the novel method on different datasets with 41,731 insect images with 9 species for training and two test datasets with 24 different taxa and 11 species. We show that the performance increases at higher ranks in the taxonomic hierarchy.

To our knowledge no other study of insect classification has proposed a hierarchical classifier with anomaly detection; however, our hierarchical classifier extends previous work where we will incorporate the hierarchy and classification in the same model. Our primary objectives are as follows:

- Classify insect species at multiple taxonomic ranks using a multitask CNN
- Determine ‘unsure’ classifications at each taxonomic rank using anomaly detection
- Evaluate model performance for species included in training data
- Test model performance for new species not included in training data

## 2. Dataset

We created four datasets to build and validated robust models that generalise well. The purpose was to demonstrate the robustness of our method to classify live insects *in situ* using automated camera monitoring or photos of manual observations. Two types of images were used: images from *in situ* monitoring with TL cameras (Bjerge et al., 2023) and images retrieved from the Global Biodiversity Information Facility (GBIF) (GBIF, 2022).

The TL images were collected prior to this work and contained observations of insects visiting *Sedum* plants. Images were annotated and validated in a process including experts as described in (Bjerge et al., 2023). The GBIF datasets derive from different kinds of sources, including museum specimens and live insects in nature taken with high-quality cameras or smartphone photos. The downloaded GBIF images were manual inspected and images were discarded containing pictures of insect larvae, images with several different species and very crowded images of insects.

### 2.1. Dataset for model evaluation

Two datasets (*TL*_*m*_ and *GBIF*_*m*_) were created for model training and validation encompassing nine insect species (Table 1). These datasets were split between training and validation with approximately 15%-20% images used for validation.

**Table 1:** Shows the nine different species of insects in the TL (*T L*_*m*_) benchmark and *GBIF*_*m*_ datasets ordered by taxonomy used for training and validation. The split of the *GBIF*_*m*_ dataset is approximately 20% for validation and 80% for training. The split of the *T L*_*m*_ dataset is approximately 16% for validation.

| Order | Family | Species | $TL_m$ | | $GBIF_m$ | |
| --- | --- | --- | --- | --- | --- | --- |
|  |  |  | Train | Validation | Train | Validation |
| Coleoptera | Coccinellidae | *Coccinella septempunctata* | 6344 | 299 | 360 | 90 |
| Diptera | Syrphidae | *Episyrphus balteatus* | 1306 | 274 | 312 | 77 |
| Diptera | Syrphidae | *Eristalis tenax* | 286 | 28 | 362 | 90 |
| Diptera | Syrphidae | *Eupeodes corollae* | 3410 | 275 | 248 | 62 |
| Hymenoptera | Apidae | *Apis mellifera* | 6934 | 1663 | 405 | 101 |
| Hymenoptera | Apidae | *Bombus lapidarius* | 1551 | 250 | 214 | 53 |
| Hymenoptera | Apidae | *Bombus terrestris* | 2955 | 240 | 396 | 98 |
| Hymenoptera | Vespidae | *Vespula vulgaris* | 956 | 201 | 396 | 98 |
| Lepidoptera | Nymphalidae | *Aglais urticae* | 286 | 27 | 97 | 24 |
|  |  | Total | 25,028 | 4,932 | 2,790 | 692 |

The TL dataset contains an imbalanced set of insect observations manually annotated with bounding boxes and classified to one of nine species (*Coccinellidae septempunctata, Episyrphus balteatus, Eupeodes corollae, Eristalis tenax, Apis mellifera, Bombus lapidarius, Bombus terrestris, Vespulas vulgaris* and *Aglais urticae*), representing the most common species in the dataset. The TL data also contain several insect detections that did not belong to these species or could not be identified with certainty. The image collection was performed on a green roof in Denmark with low variation in vegetation. While the full image resolution was large (1920 *×* 1080 pixels), the recording was done with a distance to monitor a patch of vegetation, and each insect observation covered only a small part of the full image and is therefore of low resolution. The average pixel size of the annotated bounding boxes around each insect is 70 pixels with a standard deviation of 26. The bounding boxes were cropped and re-sized to 128 *×* 128 pixels; here 98.65% of the bounding boxes, in the training dataset, are within this size.

The GBIF images (*GBIF*_*m*_) consisted of manual observations recorded with single-shot cameras of the same nine species as in the TL dataset. In these images, the insect was the focal object and they are therefore of higher resolution compared to the TL images, but the images varied in resolution and quality. Additionally, they were captured in different natural environments and contain a larger variation of background. These images were annotated with a bounding box around the individual insect species, which was cropped and re-sized to the same dimensions as the TL benchmark dataset.

### 2.2. Dataset for test with new species

Additionally, two test datasets (*TL*_*t*_ and *GBIF*_*t*_) were created to assess the model performance on species not present in the two training datasets listed in Tables 2 and 3. For *TL*_*t*_, the data consisted of detections other than those classified as one of the nine species and contains a variety of other species recorded with low frequency visiting the *Sedum* plants. These were sorted and labelled in 25 groups of taxonomy listed in Table 3. Most of the groups in this dataset are labelled at a higher taxonomy rank than the species in the train and validation dataset. The groups such as bees (Hymenoptera Apidae Apis) and hoverflies (Diptera Syrphidae) contain different species than present in the training and validation dataset. The purpose is to see how a model trained on the fully labeled taxa of species classifies these groups of *in situ* recorded insects. For *GBIF*_*t*_, the data consisted of 11 common Danish species different from the training species but within the same orders or families. These species were also chosen under the assumption that they would not be present in the *TL*_*t*_ data Table 1. The number of samples is listed in Table 2 organized in the taxonomy ranks: order, family and species. Here, genus is the first part of the word in species e.g. *Bombus* relates to bumblebees and *lapidarius* relates to the species red-tailed bumblebee.

**Table 2:** Shows the 11 different insect species in the *GBIF*_*t*_ datasets ordered by taxonomy used to test the models ability to classify new species not included in the dataset used for training.

| Order | Family | Species | $GBIF_t$<br>Test |
| --- | --- | --- | --- |
| Diptera | Syrphidae | *Eoseristalis arbustorum* | 516 |
| Diptera | Syrphidae | *Melanostoma mellina* | 421 |
| Diptera | Syrphidae | *Syrphus ribesii* | 319 |
| Hymenoptera | Andrenidae | *Andrena fulva* | 317 |
| Hymenoptera | Apidae | *Bombus hortorum* | 343 |
| Hymenoptera | Apidae | *Bombus pascuorum* | 489 |
| Hymenoptera | Megachilidae | *Megachile lagopoda* | 147 |
| Hymenoptera | Vespidae | *Vespula rufa* | 337 |
| Lepidoptera | Nymphalidae | *Aglais io* | 166 |
| Lepidoptera | Nymphalidae | *Vanessa cardui* | 350 |
| Lepidoptera | Pieridae | *Gonepteryx rhamni* | 286 |
|  |  | Total | 3,687 |

**Table 3:**
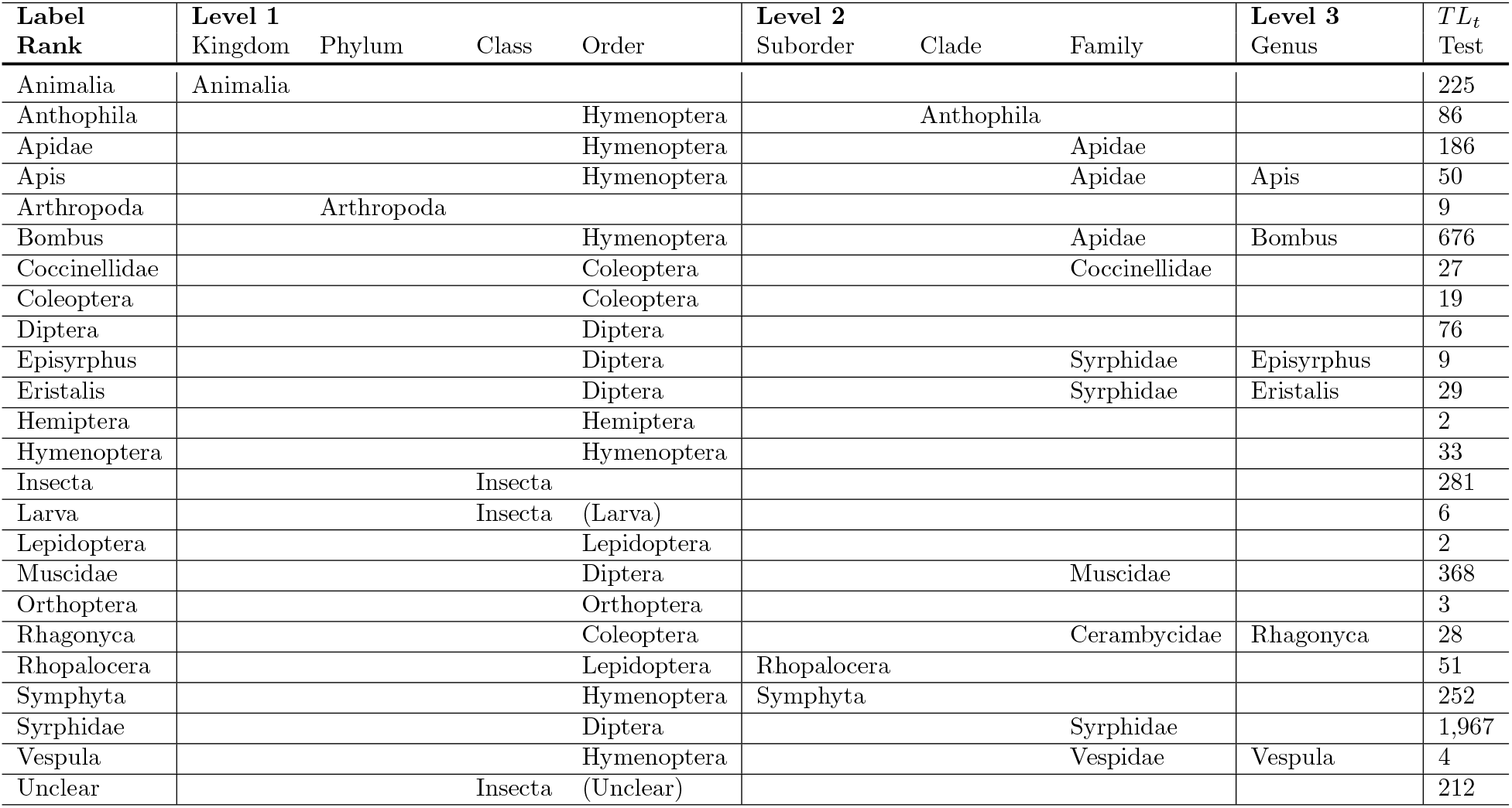
Shows the TL (*T L*_*t*_) test dataset with insect groups sorted by taxonomy rank. In total 4,601 insects and animals are labeled and sorted in 24 groups of collapsed taxa. Ranks of Kingdom, Phylum, Class and Order collapsed to level 1. Ranks of Suborder, Clade and Family collapsed to level 2. The rank of Genus belongs to level 3. The level unclear is poor quality insect images.

Some of these test species in Figure 1 are intentionally visually similar to species in the training dataset. For example, generally, only trained entomologists can tell *Bombus terrestris* and *Bombus hortum* apart from such images. We included such close species to assess if our hierarchically trained model managed to classify new species of similar and different visual appearance.

**Figure 1:**
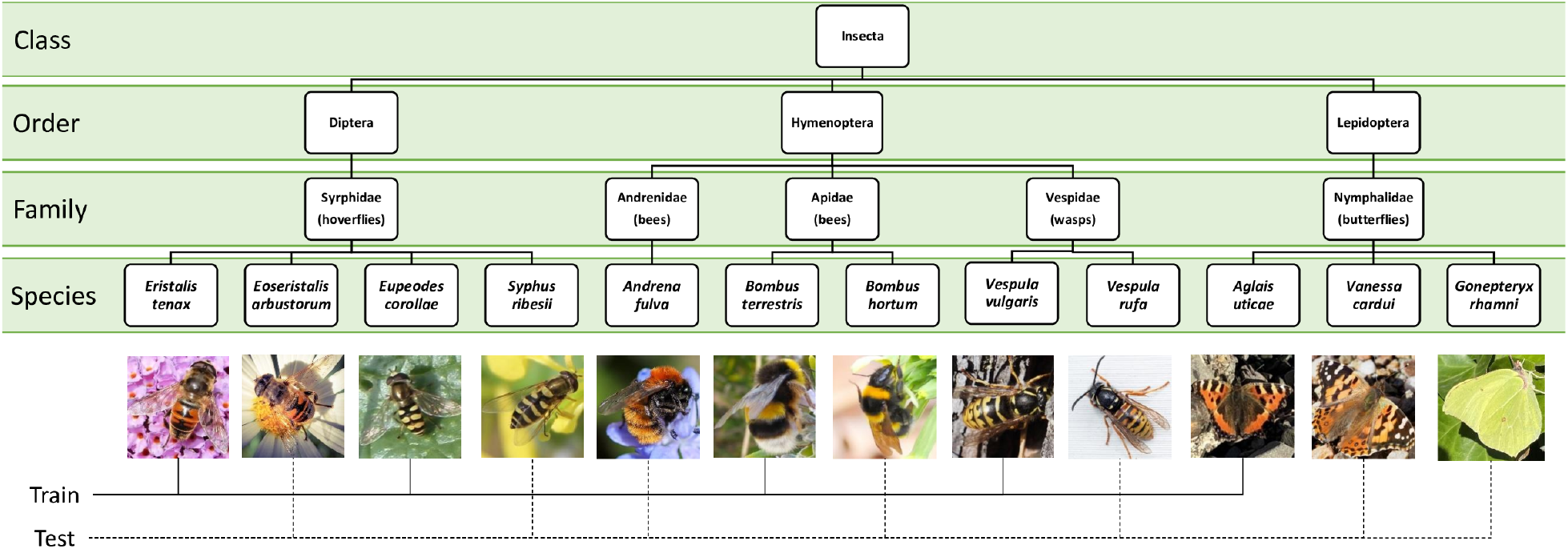
Shows an example of insect species organized in a taxonomic hierarchy used for training and testing of the classification models. Sample images of species are cropped from high-quality GBIF images. All train insect species are listed in Table 1 and test insect species are listed in Table 2. Some of the species from the train and test dataset are intentional very similar such as *Bombus terrestris* and *Bombus hortum*. The objective is to see how the trained model managed to classify new species of similar and different visual appearance.

## 3. Method

In the classification problem within deep learning, data is categorized into different classes such as sorting insect images according to species or genus. The classes are defined by the number of different labelled categories in the training data. Multi-label classification is a variant of the classification problem where multiple labels may be assigned to each insect image such as the rank of family, genus and species. In this work, we use the terminology of a hierarchy as an ordering of taxonomy ranks, where ‘levels’ are used as synonyms for ‘ranks’. The highest rank or level is numbered L1 with increased numbering (L2, L3) at lower ranks or levels. A number of classes are defined according to the number of labels in the training data defined at each rank. Classes at lower levels are nested within classes at higher levels in the hierarchy. Classes at lower levels are named ‘children’ of the associated ‘parent’ class at a higher level.

We used multitask learning (MTL) by formulating taxonomy ranks as independent tasks to learn, which we implemented by modifying the output layer of a standard CNN. Specifically, we substituted the FC layer with parallel fully connected (FC) layers for each taxonomy rank. Thus, we use hard parameter sharing between all tasks, while keeping task-specific output layers. The more rank levels that are learned simultaneously, it is assumed, the more our model has to find a representation that captures all of the taxonomic ranks and the less is the risc of overfitting on our original classification task (e.g. classification of insect species). Baxter (1997) showed that if the model has little knowledge of the true prior (few samples), then learning multiple tasks (ranks) is highly advantageous.

We simplified the network architecture and hierarchical dependency loss proposed by Gao (2020) and rewrote the code from Ugenteraan (2020), which will be explained in the following sections.

### 3.1. A multitask CNN for hierarchical classification

Our proposed network architecture is shown in Figure 2. The first part of the architecture is a CNN such as ResNet (He et al., 2016) or EfficientNet (Tan and Le, 2019). Given an input image *X* transformed by the network *N*_*CNN*_ (*θ*_*CNN*_), where *θ*_*CNN*_ is the trainable parameters of the CNN network. The network output is viewed as a root representation *R*_0_

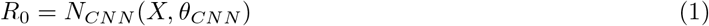

For each level *l* we add two FC layers and an activation function to perform a non-linear transformation of input *R*_0_ to level output *R*_*l*_.

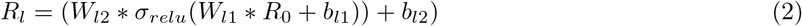

Here *W*_*l*1_ and *W*_*l*2_ represent the weights, and *b*_*l*1_, *b*_*l*2_ represents the biases for the independent FC layer representation. We apply dropout regularization in the forward pass during training so that random neurons *W*_*l*1_ are disabled in the FC branches. Each level is handled by the parallel FC layers to learn independent multiple tasks. This independent task representation is hierarchical-free and the level dependency is handled by the loss function.

**Figure 2:**
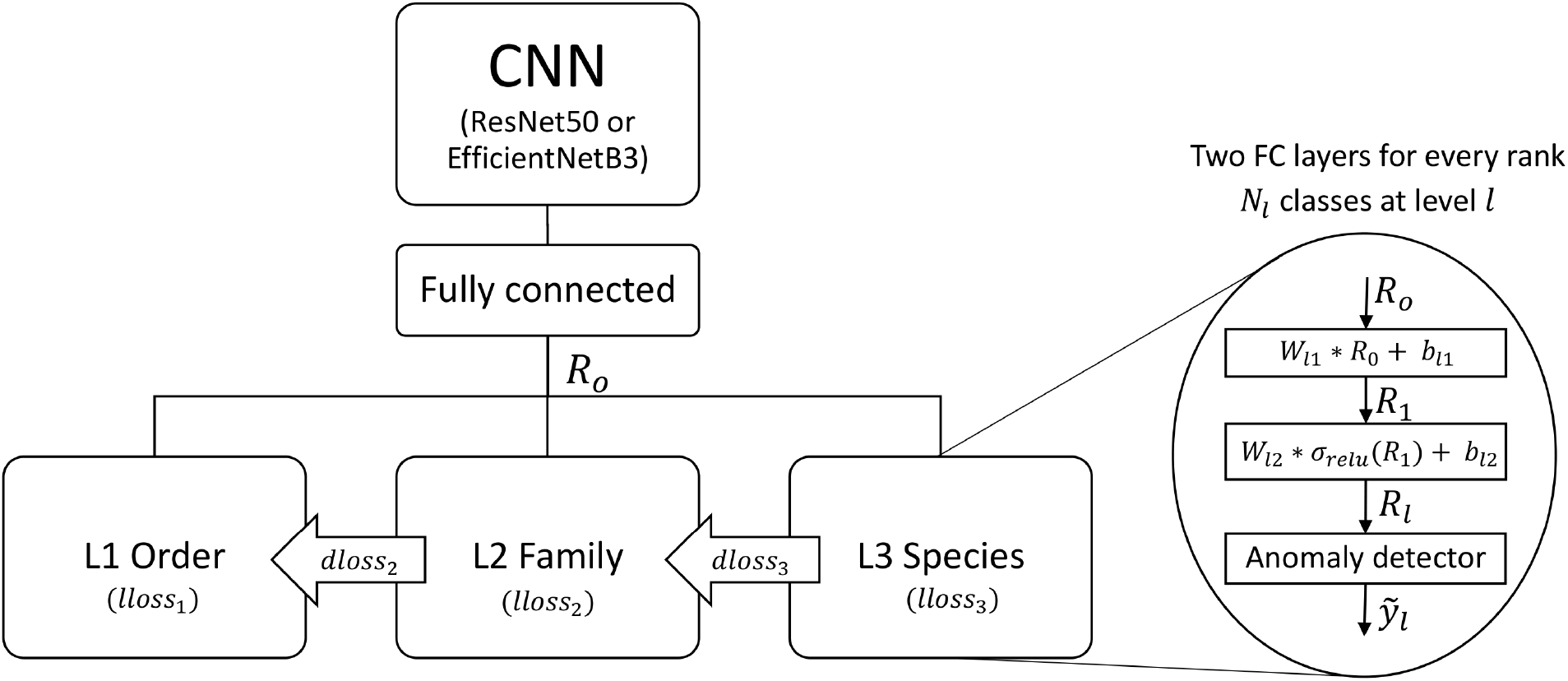
Network architecture with multitask learning to predict the three-level hierarchy. The multitask network share the same backbone (ResNet50 or EfficientNetB3) with tasks branches at the end of parallel fully connected layers. A loss function incorporates a level-dependent penalty to ensure hierarchy alignment.

#### 3.1.1. Loss function

Two types of losses are proposed by combining the cross entropy loss for each level and the new dependency hierarchical loss function. The cross entropy loss *lloss*_*l*_ (level loss) is defined as

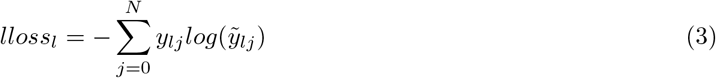

*y*_*lj*_ is the expected output of the *j*^*th*^ sample in the *l*^*th*^ hierarchy level. The hierarchical representation *R*_*l*_ Equation (2) is passed to a softmax regression layer computing the output 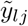. To measure the prediction errors between levels, a simplified dependence loss is defined in Equation (5). The 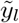 in Equation (4) denotes the predicted class in level *l*.

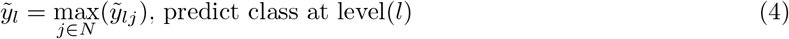

The *l*^*th*^ level dependence loss *dloss*_*l*_ is defined as

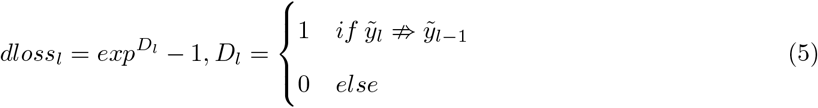

If the predicted classes of two successive levels are not parent-child relation, this dependence loss effectively penalises the learning model.

The binary *D*_*l*_ denotes whether the predicted label in the *l*^*th*^ level is a child class of the predicted class in the *l −* 1^*th*^ level. The dependence punishment 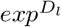 constrains the neural network to learn structure information between levels in the hierarchy. Finally, the total loss is defined as the weighted summation of the level and dependence losses, i.e.,

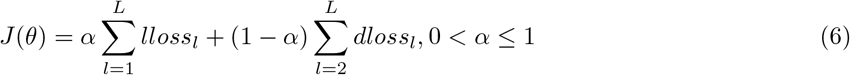

*α ∈* [0, 1] defines the priority between cross entropy level loss (*α* = 1) and dependence loss (*α* = 0). *α* must be greater than zero, but values close to zero will also work. Since we only observed marginal changes in accuracy as shown in Appendix A, we chose *α* = 0.5 for the rest of this study.

### 3.2. Anomaly detection of ‘unsure’ insect species

The concept of semi-supervised deep learning and threshold-based anomaly tagging is used to detect ‘unsure anomalies’. Often, softmax is the last layer in a classification neural network, where the maximum value determines the predicted class. Instead, we will analyze the output distribution without the softmax layer to determine outliers and predict classes. The distribution of the output *R*_*lj*_ Equation (2) for each predicted *j*^*th*^ class in the *l*^*th*^ level follows a normal distribution *R*_*lj*_ *∼* 𝒩 (*μ, σ*^2^). The output distribution is shown in Figure 3 is generated on the training dataset for correct classified inputs. The Gaussian approximation of the underlying probability density function is shown for three levels of two classes from each level. If the output value *R*_*lj*_ is below a threshold of *th* = *μ −* 2*σ*, we will predict the input as an outlier. Theoretically, an ideal normal distribution means that approximately 2.1% of correctly classified inputs will be wrongly predicted as outliers. However, when new unknown inputs are presented for the trained network, and the output lies below the threshold it will be labeled as an “Unsure” anomaly.

**Figure 3:**
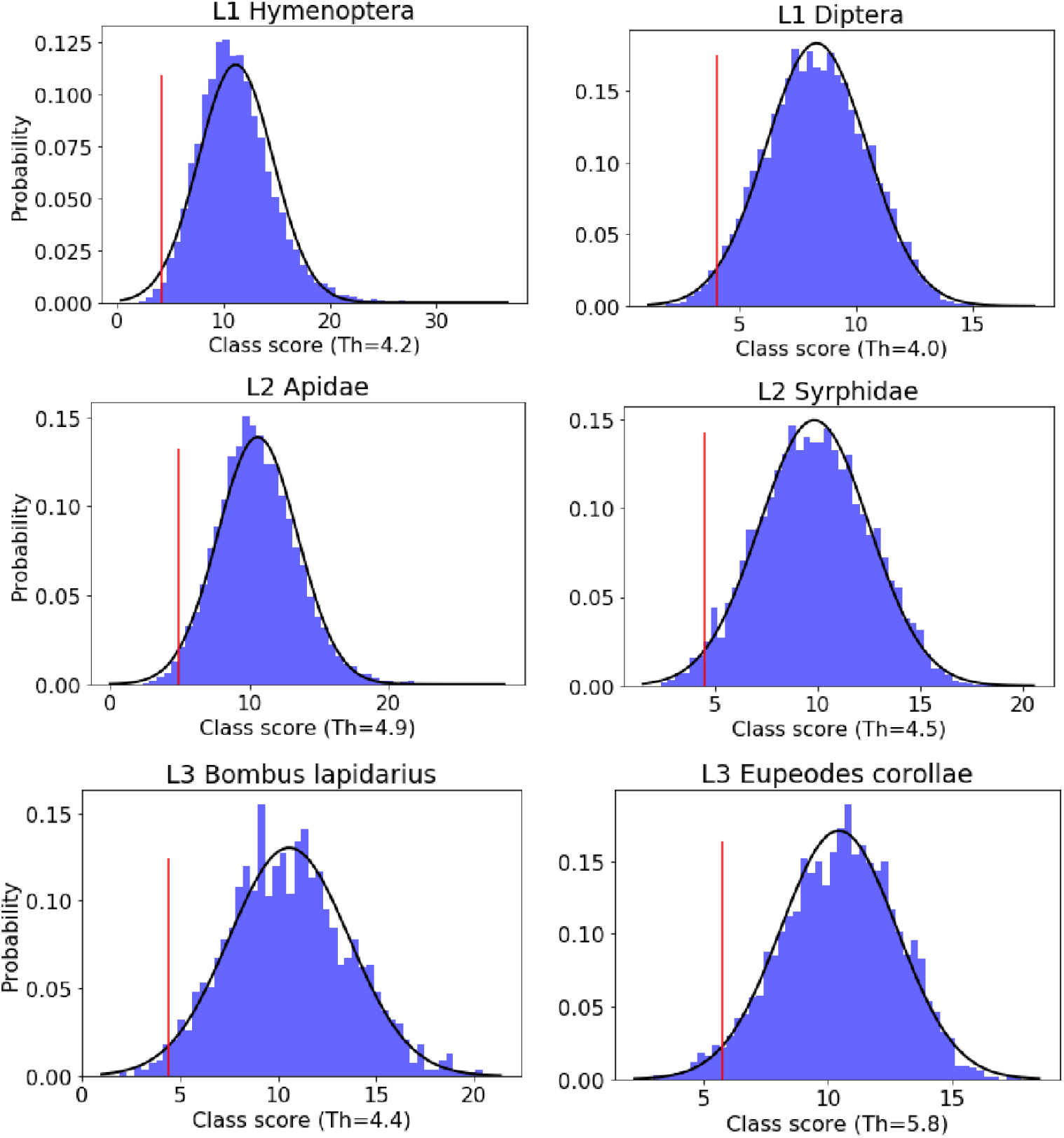
Normal approximation of the probability density function for the output *R*_*lj*_ based on the training dataset for different classes and levels in the hierarchy. The continuous lines shows the normal approximation of the observed output (histogram) of *R*_*lj*_. The red line defines the threshold (*th* = *μ −* 2*σ*) under which predictions are considered “anomalies”. and labeled “Unsure”.

Equation (4) denotes the predicted class in level *l* as the maximum output value from the network at the level. An input image is also classified as unsure when the predicted class is not correct according to the higher rank in the taxonomic hierarchy as defined in Equation (7).

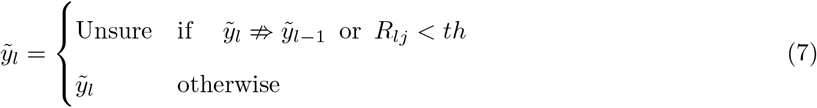

### 3.3. Training, augmentation and optimizers

Training on the datasets was performed using data augmentation including image scaling, horizontal and vertical flip, perspective distortion and adding color jitter for brightness, contrast and saturation. Data augmentation mitigates overfitting by increasing the diversity of the training data. We selected a batch size of 20 for training our models since it is faster to update, and results in less noise, than lower batch sizes. The accuracy of the models on train and validation datasets was computed after each epoch.

The Adam optimizer with a fixed learning rate of 1.0 *·* 10^*−*4^ was finally chosen after comparing with stochastic gradient descent (SGD) with the momentum of 1.0 *·* 10^*−*4^, weight decay of 1.0 *·* 10^*−*4^ and learning rate of 1.0 *·* 10^*−*3^. SGD was tested with the parameters as recommended by L. Smith (Smith, 2018), but achieved a 1–2% decrease in accuracy compared to using the Adam optimizer.

The ResNet50 architecture (He et al., 2016) was modified and trained with transfer learning using pretrained weights from ImageNet (Smith, 2018). Models trained with transfer learning outperformed models trained with random weights with an increase of 4% in F1-Score at the species level. To compare with other state-of-the-art networks EfficientNetB3 (Tan and Le, 2019) was also evaluated with pre-trained weights from ImageNet. Transfer learning involves training a model on data from a source domain *T*_*S*_ = *P* (*y*|*X*) and then transferring it to a target domain *T*_*T*_ = *P* (*y*|*X*) — typically with less data available. In this case, ImageNet contains 1,000 classes with 1,281,167 images for training and 50,000 for testing, while the train insect TL (*TL*_*m*_) benchmark dataset only contains 9 classes with 25,028 TL images.

First, we used the Cifar-100 (Krizhevsky et al., 2009) dataset as a quality control to show we can reproduce the same performance as achieved by Gao (2020). We performed a test on the Cifar-100 dataset with 28 *×* 28 pixel images with a two-level hierarchy of 20 super-classes and 80 subclasses with a total of 500 training and 100 validation images per class without pre-trained weights. Our MTL model with a simplified loss function Equation (5) performed very similarly to the original model presented by Gao (2020) with code from Ugenteraan (2020). We achieved an accuracy of 81.64% for super-classes and 71.61% for subclasses Gao et al.’s model – compared to 81.83% for super-classes and 70.93% for subclasses with our MTL model.

## 4. Results and discussion

In this section, different models were trained and evaluated on the TL and GBIF datasets. First, models were trained on time-lapse (*TL*_*m*_) images to evaluate the hierarchical classification with ResNet and EfficientNet with MTL using transfer learning described in section 4.1. The best model was then evaluated on the test dataset (*TL*_*t*_) to see how new species were handled by the hierarchical classifier with anomaly detection described in section 4.2. In section 4.3 additional two models were trained with ResNet50 on the GBIF dataset (*GBIF*_*m*_) and a mixture (MIX) of the *TL*_*m*_ and *GBIF*_*m*_ datasets. These models were tested to assess how different trained models perform on the TL and GBIF dataset. The model trained on the MIX dataset was finally evaluated on the GBIF test dataset (*GBIF*_*t*_) to assess how new species were classified in the taxonomic hierarchy or categorized as unsure insects by the anomaly detector.

### 4.1. Model evaluation on TL images

In this section during the model evaluation of the CNN classifiers, the anomaly detection filer was not used. Models with ResNet50 and EfficientNetB3 were trained on the TL (*TL*_*m*_) benchmark dataset with transfer learning. Choosing the best model based on minimum total loss during model training is further described in Appendix B.

Five models were trained for each combination of models (ResNet50 and EfficientNetB3) with and without hierarchical classification (MTL) with a total of 20 models. Level-3 accuracy *∈ {*98.0, 97.8, **98.3**, 97.7, 97.9*}* for the five models trained with ResNet50 and MTL. Here, the best-performing ResNet50MTL model was later used to evaluate the hierarchical classification on the validation (*TL*_*m*_) and test (*TL*_*t*_) dataset.

The average and standard deviation of precision, recall and F1-score are shown in Table 4 for the ResNet50 and EfficientNetB3 models validated with multitask learning and pre-trained weights. As expected, we observe higher precision, recall and F1-score at higher taxonomic ranks. Here we see an increase in average F1-score ranging from 95.7% (L3) to 98.7% (L2) and 99.1% (L1) at the highest ranks with ResNet50MTL. A resent study by Ong and Hamid (2022) designed to classify five taxa of museum insect species achieved Fl-scores below 90% with separate models for each level of the hierarchy. However, our dataset is more comprehensive with a lower image resolution.

**Table 4:** Shows the average performance and standard deviation calculated on five trained models. It shows the average precision, recall and F1-score for trained ResNet50 and EfficientNetB3 (EffNetB3) models modified for multitask learning (MTL) with transfer learning using pre-trained weights from ImageNet. The models are trained and validated on the TL dataset *T L*_*m*_. The models ResNet50, EfficientNetB3 are trained without MTL.

| Model | Level | Precision |  | Recall |  | F1-score |  |
| --- | --- | --- | --- | --- | --- | --- | --- |
| | | Avg | Std ($10^{-3}$) | Avg | Std ($10^{-3}$) | Avg | Std ($10^{-3}$) |
| ResNet50MTL | L1 Order | 0.990 | (1.0) | 0.991 | (1.1) | 0.991 | (0.5) |
| EffNetB3MTL | L1 Order | 0.986 | (4.4) | 0.993 | (0.5) | 0.989 | (2.3) |
| ResNet50MTL | L2 Family | 0.987 | (0.8) | 0.986 | (0.9) | 0.987 | (0.7) |
| EffNetB3MTL | L2 Family | 0.984 | (3.1) | 0.988 | (0.7) | 0.986 | (1.6) |
| ResNet50MTL | L3 Species | 0.955 | (4.3) | 0.961 | (9.8) | 0.957 | (3.2) |
| EffNetB3MTL | L3 Species | 0.948 | (5.2) | 0.966 | (5.1) | 0.956 | (1.7) |
| ResNet50 | Species | 0.955 | (3.3) | 0.957 | (7.3) | 0.955 | (0.4) |
| EffNetB3 | Species | 0.953 | (2.5) | 0.966 | (2.5) | 0.959 | (0.8) |

The performance of ResNet50 and EfficientNetB3 with transfer learning is very similar; however, at the species level (3), ResNet50 has a higher standard deviation (0.32%). State-of-the-art models trained with transfer learning and without MTL have an F1-score of 96% similar to models trained with MTL at the species level. The models trained with ResNet and EfficientNet have the absolute best performance and outperform the benchmark YOLOv5 model on the TL dataset referenced in paper (Bjerge et al., 2023). This is expected since the tasks performed by YOLOv5 include both object detection as well as classification. No other state-of-the-art CNN models were evaluated since it is not expected to get higher scores than in the area of 96% to 99% and the purpose was to demonstrate that different CNNs can be modified for hierarchical multitask learning.

Figure 4a shows the hierarchical structure with accuracy for classification at each level of the taxonomy (Best ResNet50MTL model). Parents with more children, in most cases, show higher accuracy than child nodes. The accuracy was in general very high for all classes, with many close to 100%. Some closely related species were misclassified; for example, 1.3% of *Bombus terrestris* were classified as *Bombus lapidaries*. Species in the family Syrphidae were similarly misclassified. Note that 11% of the species *Eristalis tenax* were classified as *Apis mellifera. Eristalis tenax* is superficially similar to, and widely considered to be a Batesian mimic of, *Apis mellifera* (Golding et al., 2005).

**Figure 4:**
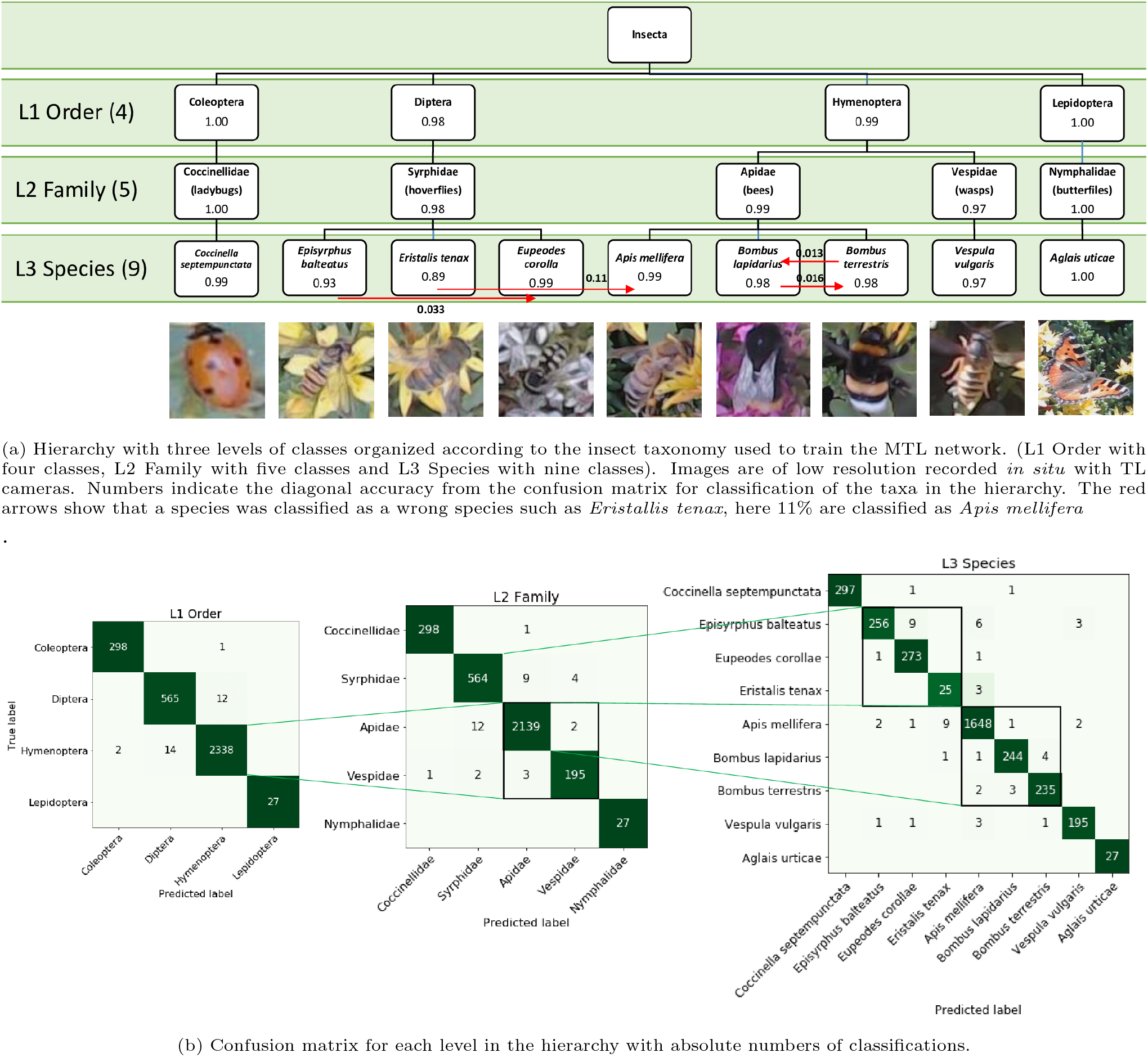
Validation of insects in taxonomic hierarchy

The confusion matrix Figure 4b shows for each level of the hierarchy the absolute numbers of classifications, of which very few were incorrect. The high numbers in the diagonal of the confusion matrix indicate correct classifications. At the order level, only 14+12 insects from the orders Diptera and Hymenoptera were wrongly classified. At the family level, there were 12+9 insects from the families Syrphidae and Apidae that were wrongly classified. At the species level, it was obvious that *Eristalis tenax* and *Apis mellifera* contribute most (9+3) of the misclassifications. The class imbalance, with few samples of *Eristalis tenax*, probably contributes to these errors. In total, 57 wrong predictions were found at the species level, where 8 predictions did not fit the taxonomy. Some predictions did not fit the taxonomy (i.e. there was a mismatch between predicted order, family, and/or species). For example, the species *Apis mellifera* does not belong to the matching predicted families Coccinellidae and Syrphidae and the family Vespidae does not belong to orders Coleoptera and Diptera. Some examples are listed below (misfits in bold):

- Coleoptera, Coccinellidae, ***Apis mellifera***
- Coleoptera, Coccinellidae, ***Eupeodes corollae***
- **Coleoptera**, Vespidae, *Vespula vulgaris*
- Diptera, Syrphidae, ***Apis mellifera***
- Diptera, **Vespidae**, *Episyrphus balteatus*
- Hymenoptera, Apidae, ***Eristalis tenax***

Theses results shows that only few predictions did not fit the taxonomy and they would be classified as unsure when the anomaly detector is applied in the next section.

### 4.2. Test with new species in TL images

We assessed the ability of models to (1) classify unfamiliar species to higher taxonomic ranks and (2) highlight where a reliable classification could not be reached. To do so, we evaluated the best model (ResNet50MTL) against a TL test dataset (*TL*_*t*_) containing specimens that could only be identified as “unsure” or to the level of order, family or genus. Anomaly detection was used in this evaluation of the results. Table 5 shows an increase in precision and F1-score at higher taxonomic ranks where recall is more steady. Figure 5 shows the confusion matrix when classifying insects at each taxonomic rank. Out of 4,601 samples, 126 (2.7%) had predictions that did not fit the taxonomy. These were added to the “Unsure” predicted labels.

**Table 5:** Shows the precision, recall and F1-score on the TL test dataset (*T L*_*t*_) at each taxonomic rank. Unclear annotated species are not included in these metric calculations.

| Level | Precision | Recall | F1-score |
| --- | --- | --- | --- |
| L1 Order | 0.766 | 0.902 | 0.828 |
| L2 Family | 0.708 | 0.910 | 0.796 |
| L3 Species | 0.304 | 0.877 | 0.452 |

**Figure 5:**
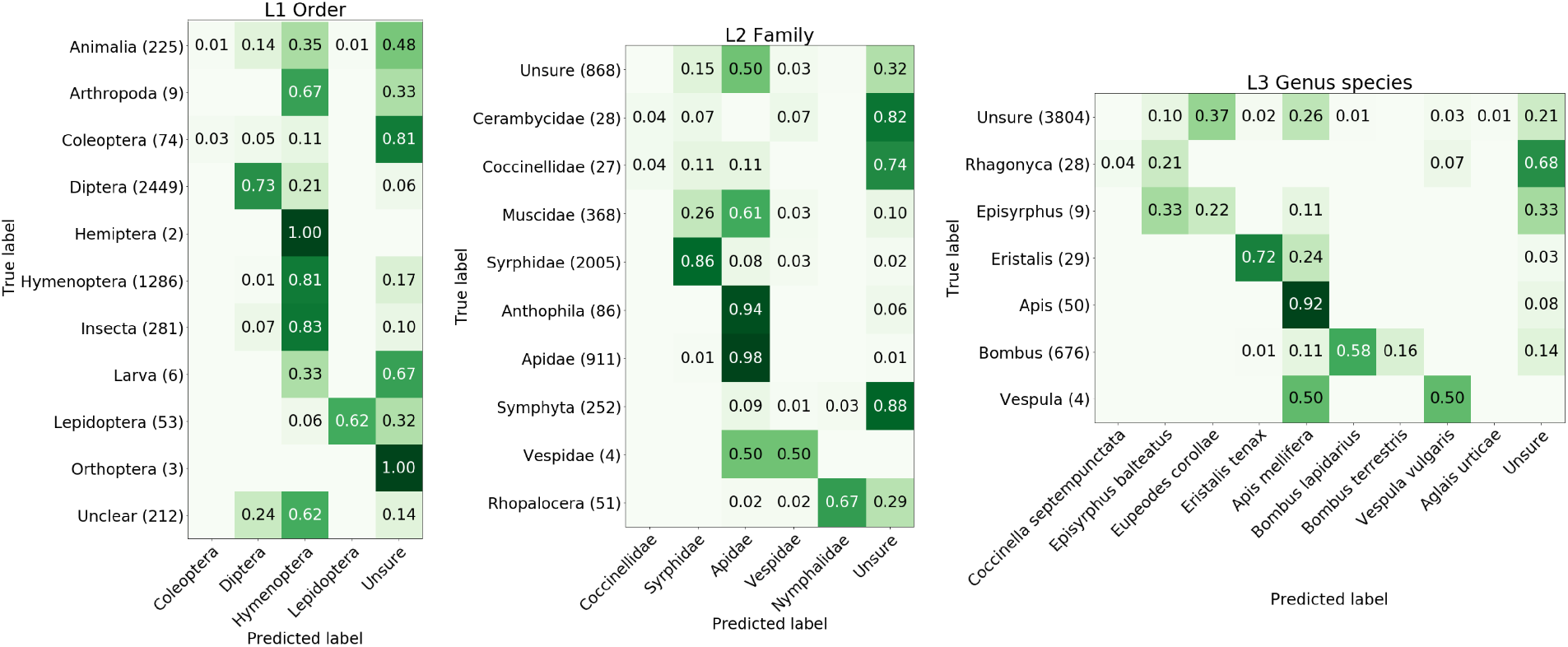
Confusion matrix for performance on the test dataset (*T L*_*t*_). Unsure true labels mean that the group of insects was not identified and labelled at this level of the taxonomy. The unsure predicted label indicates a classification filtered by the anomaly detector.

At the order level, 81% of Hymenoptera, but only 73% of Diptera, were correctly classified. Image ground truth assigned to the Kingdom of Animalia, Class of Insecta and Larva were mostly classified as Hymenoptera. Unclear insects of poor quality images were mostly predicted as Diptera and Hymenoptera and 14% were predicted as unsure insects. For the order Coleoptera, 81% were classified as unsure, probably because many specimens of the family Conccinellidae did have a visually different appearance than *C. septempunctata* represented in the training data. At the family level, 98% of Apidae and 86% of Syrphidae were correctly predicted. Anthophia were 94% correctly classified as bees (Apidea).

Only 10% of the family of houseflies (Muscidae) were predicted as unsure insects and most (61%) were predicted as bees. At the genus level, most (74%) of the Bombus were predicted as either *Bombus lapidarius* or *Bombus terrestris* as we would expect. Images in the genus Rhagonyca were 68% classified as unsure insects. This is probably due to a visually different appearance than the species included in the training dataset (*TL*_*m*_).

The general findings are that the rank of order and family will typically be predicted as the order Hymenoptera or family Apidae. This is probably because these insects are most dominant in the imbalanced training dataset *TL*_*m*_. New insect families, with different appearances present in the training data, such as Cerabycidae and Symphyta and the genus Rhagonyca are correctly predicted as unsure predictions (anomalies).

### 4.3. Test with new species in GBIF images

To classify species in the *GBIF*_*t*_ test dataset we found that models need to be trained with GBIF data or a mixture of TL and GBIF images to gain high performance. Therefore, two ResNet50 models were trained one on the *GBIF*_*m*_ dataset and another on a mixture (MIX) of the TL (*TL*_*m*_) and *GBIF*_*m*_ images. The models trained and validated on the datasets (*GBIF*_*m*_ and MIX) performed very similarly to the model trained on the *TL*_*m*_ dataset with an F1-score above 96% on all three levels. However, a model trained on either *GBIF*_*m*_ or *TL*_*m*_ data performed poorly on the opposite validation dataset with an F1-score of 68% at species level 3, see Table 7 and Figure 7. On family level 2 and order level 1, the F1-score was 70% for a trained TL model validated on the *GBIF*_*m*_ dataset. For a trained GBIF model validated on the *TL*_*m*_ dataset, the F1-score was higher with 77% at level 2 and 83% at level 1. For the model trained on a MIX dataset, the F1-score raised to 96% (L3), 98% (L2) and 99% (L1) which are similar for models only trained on the *GBIF*_*m*_ or *TL*_*m*_ dataset. The trained model on the MIX dataset has a similar high performance on the *TL*_*m*_ and *GBIF*_*m*_ validation datasets, see Table 7. This indicates a more robust model being able to classify images from both TL images and photos of manual observations.

These results show that F1-scores increase at higher levels in the hierarchy of taxonomy rank, especially for the models validated on opposite datasets in Figure 7. They also show that it is important with a high variation in the images of insect species to train a robust classifier. It seems not to be sufficient with only GBIF images, even though the variation of image resolution, quality, and background vegetation is high.

A final test was performed using the *GBIF*_*t*_ test dataset to see how a trained model performed on images with new species. The ResNet50 model trained on the MIX dataset evaluated on the *GBIF*_*t*_ test dataset is shown in Figure 6. Table 6 shows the same pattern as test on the *TL*_*t*_ dataset with an increase in precision and F1-score at higher taxonomic ranks. The recall is 1.00 at the species level since none of the *GBIF*_*t*_ test species were present in the trained model.

**Table 6:** Shows the precision, recall and F1-score on the GBIF test dataset (*GBIF*_*t*_) at each taxonomic rank.

| Level | Precision | Recall | F1-score |
| --- | --- | --- | --- |
| L1 Order | 0.904 | 0.803 | 0.850 |
| L2 Family | 0.756 | 0.822 | 0.788 |
| L3 Species | 0.331 | 1.000 | 0.498 |

**Figure 6:**
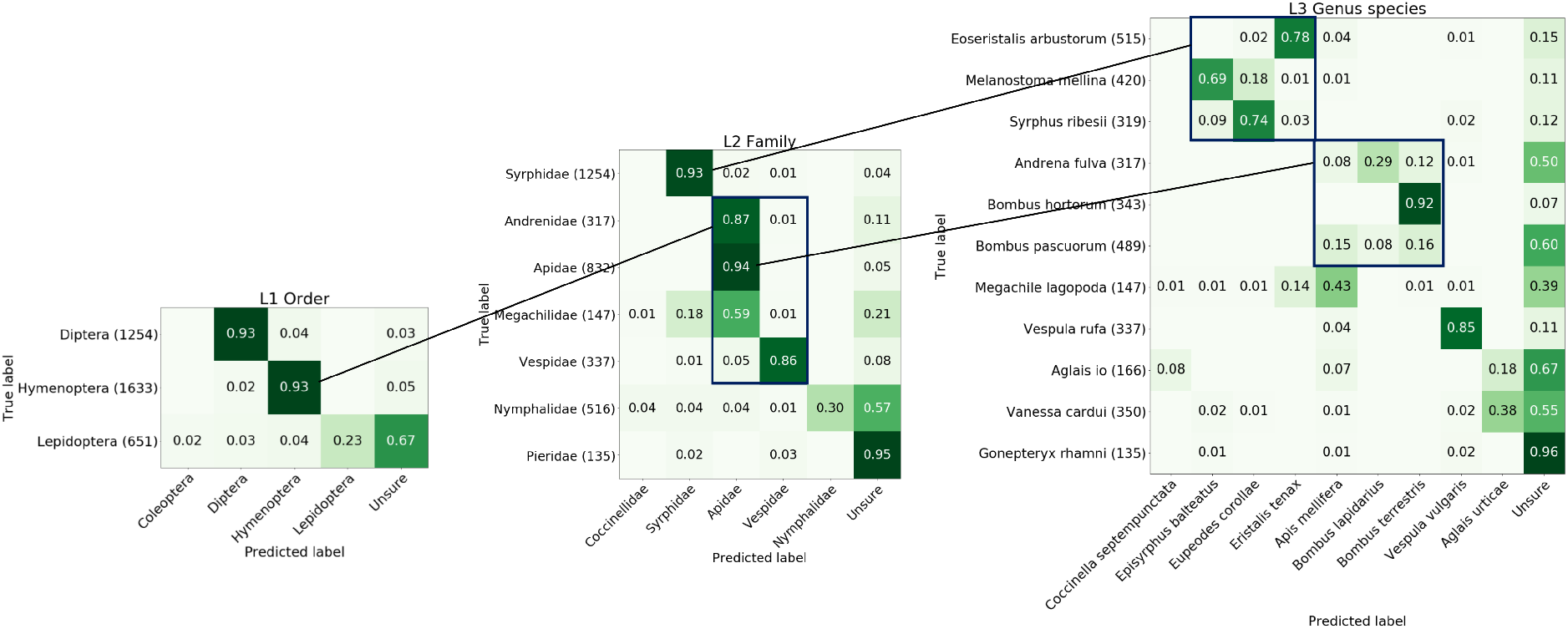
Confusion matrix for the *GBIF*_*t*_ test dataset of new species in the taxonomic hierarchy.

**Figure 7:**
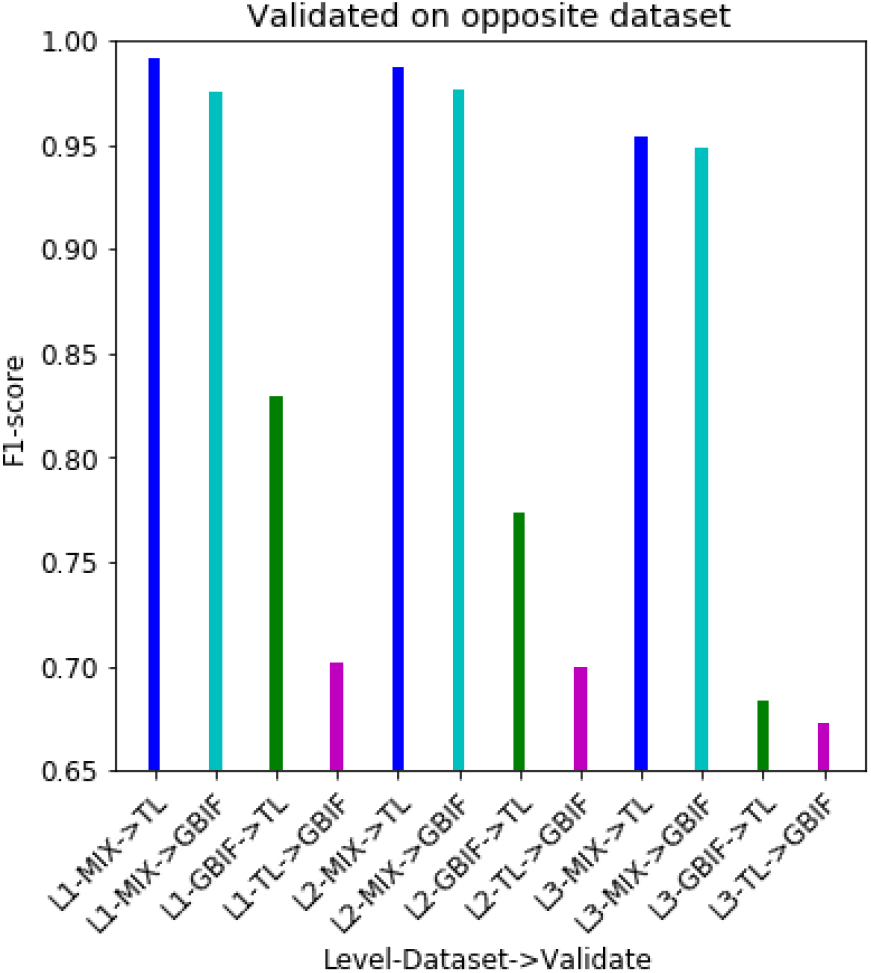
F1-scores for each level in the hierarchy of models trained on three different datasets (*GBIF*_*m*_, *T L*_*m*_ or MIX). “Validation on opposite dataset” shows how models perform on the *GBIF*_*m*_ or *T L*_*m*_ validation datasets listed in Table 7.

**Table 7:** Shows the F1-scores for each level in the hierarchy of models (MIX, GBIF and TL) trained on the three different datasets (*GBIF*_*m*_, *T L*_*m*_ and MIX) and validated on either the *T L*_*m*_ or *GBIF*_*m*_ validation dataset.

| Level-Model | Validated | F1-score |
| --- | --- | --- |
| L1-MIX | TL | 0.991 |
| L1-MIX | GBIF | 0.975 |
| L1-GBIF | TL | 0.829 |
| L1-TL | GBIF | 0.702 |
| L2-MIX | TL | 0.987 |
| L2-MIX | GBIF | 0.976 |
| L2-GBIF | TL | 0.774 |
| L2-TL | GBIF | 0.700 |
| L3-MIX | TL | 0.954 |
| L3-MIX | GBIF | 0.948 |
| L3-GBIF | TL | 0.683 |
| L3-TL | GBIF | 0.673 |

At level 1, the order Diptera and Hymenoptera were classified correctly with an accuracy of 93%. 67% of the insects of the order Lepidoptera were classified as unsure; this was due to only one species (*Aglais urticae*) being included in the training data.

At level 2, the family Syrphidae and Vespidae were classified correctly with an accuracy of 93% and 86%. Bees in the clade Anthophila of the family Andrenidae, Apidae and Megachildae were classified as Apidae with an accuracy of 87%, 94% and 59%. Only 30% of Nymphalidae were correctly classified, this was due to an insufficient number of different species in this category.

At level 3, some species shown in Figure 1, were classified as having the most similar appearance of species in the training dataset. Only species (*Andrena fulva, Bombus pascuorum, Aglais io*, and *Gonepteryx rhamni*) with a very different appearance were classified correctly as unsure. This was especially the case for Common brimstone (*Gonepteryx rhamni*) of color citrus here 96% was classified correctly as unsure.

### 4.4. Anomalies classified correct at higher rank

If an unsure prediction (Anomaly) is correct at a higher level of the taxonomy the classification would be valid and useful. Table 8 shows the number of unsure predictions of correct classification at higher taxonomy rank on the two test datasets. At L2 45.7% and 77.6% of unsure family predictions were correctly classified at the order rank. At L3 32% and 67.4% of unsure species predictions were correctly classified at the family rank.

**Table 8:**
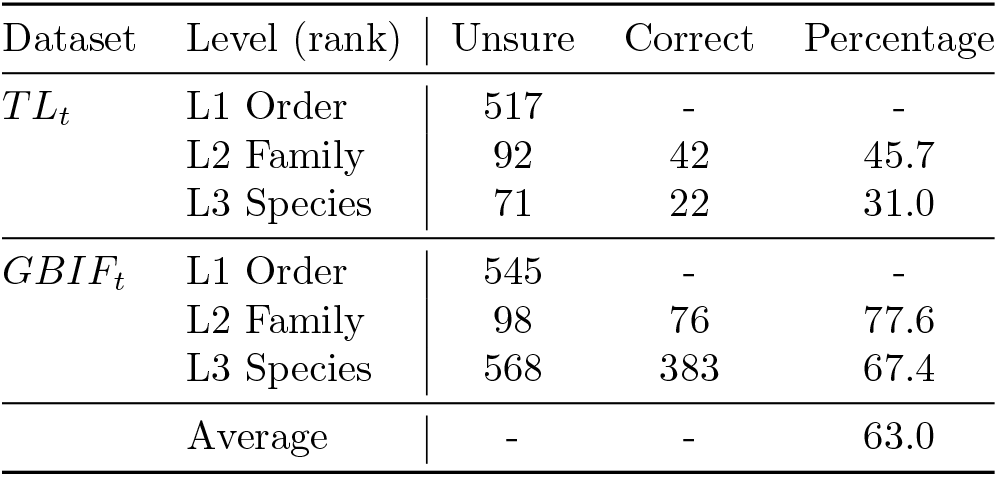
Shows the number of correct classification at higher taxonomy rank for the TL (*T L*_*t*_) and *GBIF*_*t*_ test dataset. Unsure: the number of unsure prediction if the higher rank has a correct prediction. Correct: the higher rank is correct classified for an unsure prediction at level Lx. Percentage: relative number of correct predictions at higher rank.

In total 565 unsure species have been correctly classified at higher taxonomy rank, which is 63% of all unsure anomalies. This result indicates that the anomaly detector does contribute to filter species, especially of visually different appearance such as *Gonepteryx rhamni, Aglais io, Bombus pascuorum* and *Vanessa cardui* with unsure classifications above 50% as shown in Figure 6.

### 4.5. Potential applications

Our method has direct applications for image-based monitoring of insects in natural environments, both for agriculture and biodiversity conservation (Ratnayake et al., 2021; Bjerge et al., 2021a; Preti et al., 2021). One could combine our presented hierarchical classifier with an object detector such as from the family of real-time YOLO detectors (Terven and Cordova-Esparza, 2023). The object detector could be trained as a generic insect detector by creating a general dataset with one universal “insect” class based on the annotations in the *TL*_*m*_ dataset (Bjerge et al., 2023). Ideally, this would be supplemented with annotated images of a variety of background vegetation with insects to ensure a more robust detector. For time-lapse detection, a YOLO model with motion enhancement could improve the detection of insects as suggested by (Bjerge et al., 2022). Subsequent to detection, insects could be classified into taxonomic ranks as proposed in this paper. In contrast to the universal insect detector, the hierarchical classifier should be trained on taxa relevant to the geographic and temporal scope of the setting. Here, GBIF data could provide training data, supplemented with insect image data from the camera location to achieve high accuracy. Since it would be infeasible to get sufficient samples of all potential species, the anomaly detector is relevant for flagging unseen and unsure species. This two-step approach would be more comprehensive and flexible than training an object detector to both detect and classify insect species within a single model.

Hierarchies are pervasive across deep learning applications. However, hierarchy is especially relevant when classifying living things, as each species has a unique position on the tree of life. Insects are particularly diverse, comprising a great deal of important pollinator, pest, prey and decomposer species. Furthermore, many insect species cannot be verified without dissection or genetic analysis. By leveraging known relationships between easily identifiable species, our approach helps to classify unverifiable specimens to a higher taxonomic level. This could be especially useful in pest detection applications, for example where cryptic relatives of pest species must also be detected (Preti et al., 2021).

Novelty and uncertainty are also pervasive issues in deep learning. By combining anomaly detection with multitask transfer learning, we show where a specimen likely belongs to a novel species, but also indicate a likely higher taxonomic rank and class. One application of this could be to notify relevant experts about novel insect specimens, allowing efficient discovery of new species or colonization events. Another application is to better represent uncertainty within automated insect monitoring systems. Downstream users of the data can then choose to throw out novel or uncertain classifications or analyze their counts at a higher taxonomic level.

## 5. Conclusion

We have demonstrated the possibility of hierarchical insect classification based on taxonomic ranks. The accuracy of our CNN increased at higher ranks and achieved F1-scores in the range of 95% to 99%.

The modified ResNet50 and EfficientNetB3 with hierarchical multitask learning performs similarly to a state-of-the-art network for species classification. However, by introducing hierarchical models, we correctly classified 63% of anomalies at higher taxonomic levels. State-of-the-art networks with “flat” classifiers will not correctly classify anomalies but assign unfamiliar species to insect classes in the train datasets. The presented method for anomaly detection performs best when novel species are visually different from known species in the training dataset.

New insect species in families or orders included in the trained model were classified correctly such as the orders Diptera and Hymenoptera as well as the families Syrphidae (hoverflies) and Apidae (bees). Insects of a new family such as houseflies (Muscidae) were predicted as bees (Apidae) which was the highest-frequency family in the training data. That means a model trained with an imbalanced dataset will likely detect new taxa of insects to a similar class with most data samples.

Supplementing models with GBIF data makes the classifier more general and robust to classify images from both TL and photos of manual observations. The model trained on a mixture of TL and GBIF images predicts taxa ranks correctly on a GBIF test dataset with new insect species. Insects were classified as species most similar in appearance and species with a very different appearance were classified correctly as an unsure anomaly.

Our work represents a valuable method for future applications in entomology for automated insect classification using TL recordings or manual observations with captured images of live insects.

## Declaration of Competing Interest

None.

## Acknowledgments

The work was supported by the European Union’s Horizon Europe Research and Innovation programme, under Grant Agreement no. 101060639 (MAMBO).

---

**Figure A.1:**
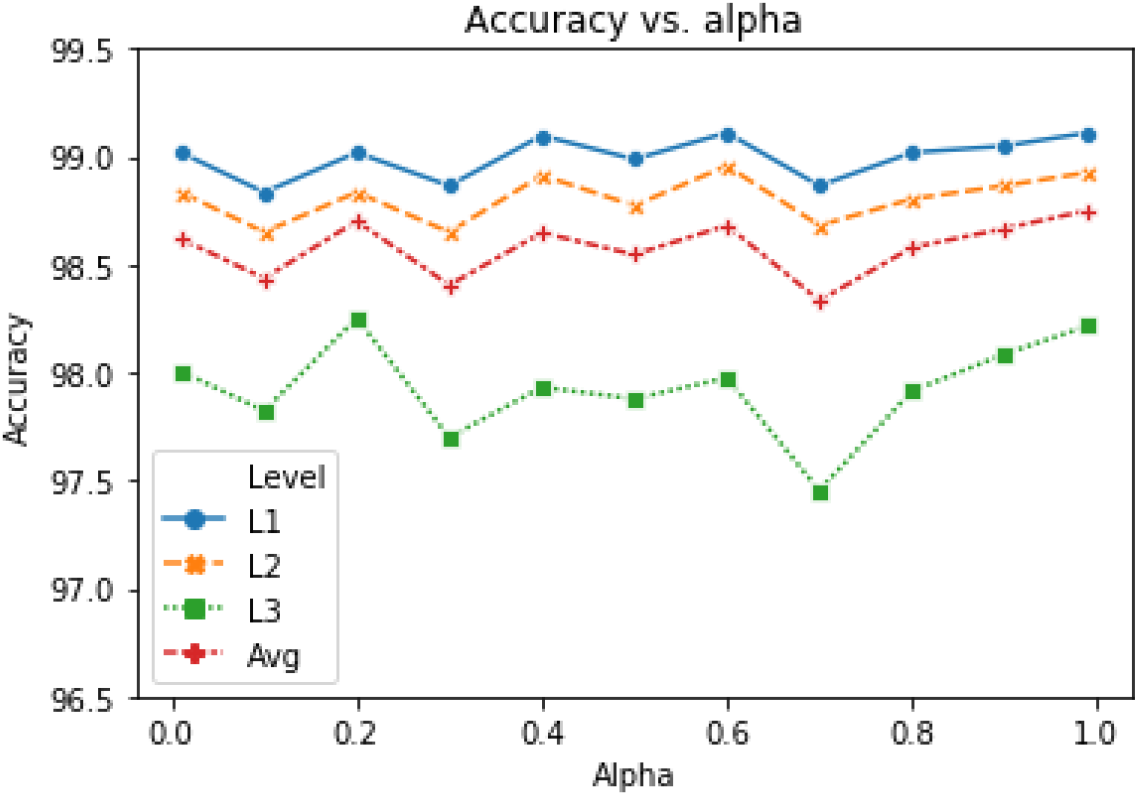
Validation accuracy for each level in the hierarchy as a function of the *α* value used in the total loss function. For level 3 the largest variation in accuracy is only 0.80% and the average variation is less than 0.42%.

## Appendix A

### Evaluation of the alpha parameter

We trained several models on the insect TL (*TL*_*m*_) benchmark dataset to optimise the hyper-parameter *α*, with *α ∈* [1.0 *·* 10^*−*6^, 1.0] achieving high accuracy as shown in Figure A.1. Since we only observed marginal changes in accuracy, for all levels, we chose *α* = 0.5 in our study.

## Appendix B

### Training of models

The best model with transfer learning was chosen based on the minimum total loss after nine epochs as seen in Figure B.2. Note that we observe overfitting after nine epochs where the validation loss starts to increase, although the bias is still very low. The increase is indicated by a higher difference between train and validation loss and bias is the loss evaluated on the training dataset.

Note that the largest variation is 0.6%, which is very similar to the variation of 0.8% when training with different *α* values in Figure A.1. This indicates a minimal impact in change of accuracy for different choices of the *α* value.

**Figure B.2:**
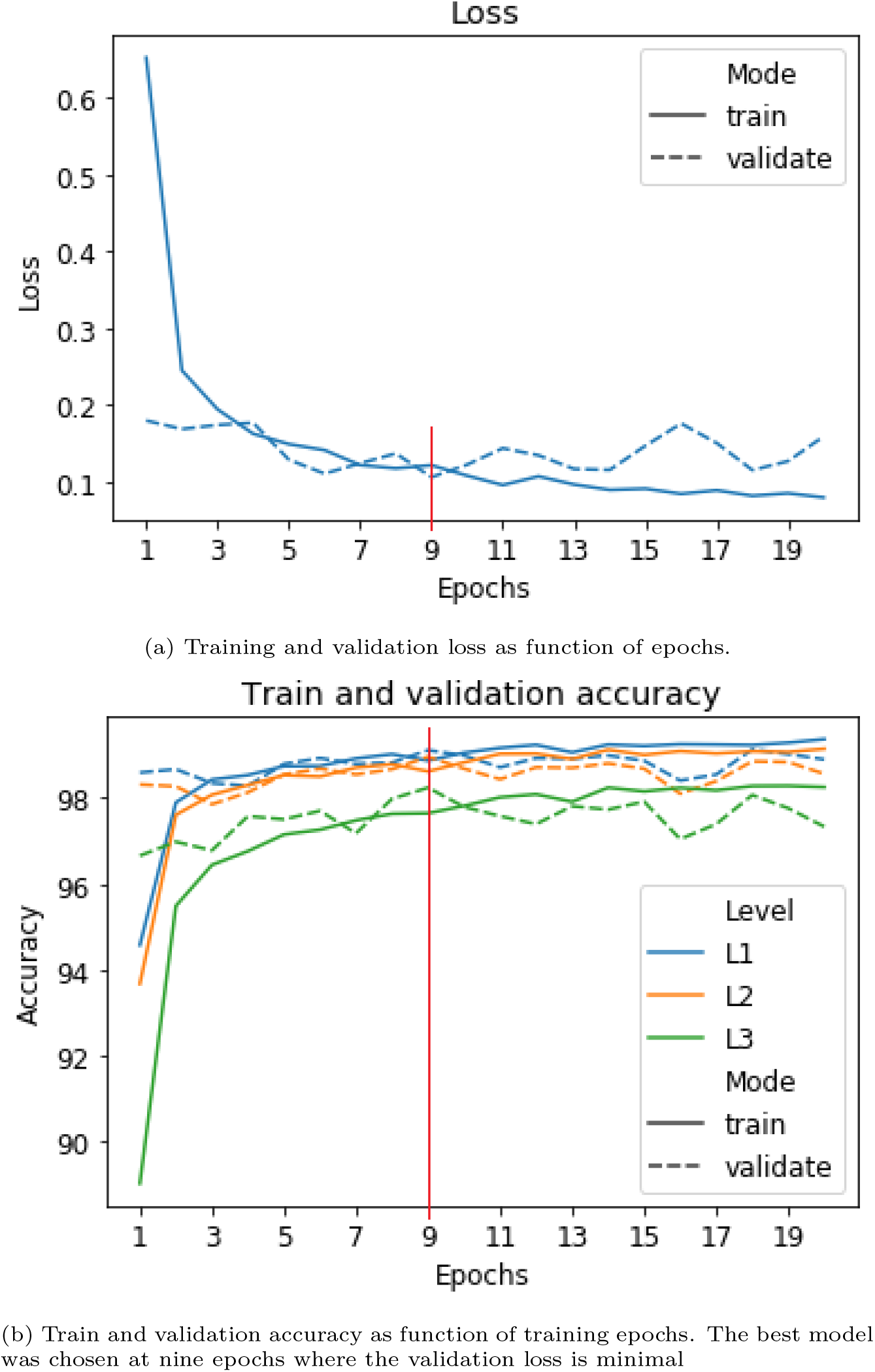
Loss and accuracy during training of the ResNet50 MTL model with transfer learning.

## Notes

https://vision.eng.au.dk/ecostack/

### Competing Interest Statement

The authors have declared no competing interest.

## References

An G, Akiba M, Omodaka K, Nakazawa T, Yokota H. Hierarchical deep learning models using transfer learning for disease detection and classification based on small number of medical images. Scientific Reports 2021;11(1). doi:10.1038/s41598-021-83503-7.

Baxter J. A Bayesian/Information Theoretic Model of Learning to Learn via Multiple Task Sampling. Machine Learning 1997;28(1). doi:10.1023/A:1007327622663.

Bertinetto L, Mueller R, Tertikas K, Samangooei S, Lord NA. Making better mistakes: Leveraging class hierarchies with deep networks. Proceedings of the IEEE Computer Society Conference on Computer Vision and Pattern Recognition 2020;:12503–12 doi:10.1109/CVPR42600.2020.01252. arXiv:1912.09393.

Bjerge K, Alison J, Dyrmann M, Frigaard CE, Mann HMR, Høye TT. Accurate detection and identification of insects from camera trap images with deep learning. PLOS Sustainability and Transformation 2023;2(3):1–18. doi:10.1371/journal.pstr.0000051.

Bjerge K, Frigaard CE, Karstoft H. Motion informed object detection of small insects in time-lapse camera recordings. arXiv 2022;arXiv:2212.00423.

Bjerge K, Mann HM, Høye TT. Real-time insect tracking and monitoring with computer vision and deep learning. Remote Sensing in Ecology and Conservation 2021a;doi:10.1002/rse2.245.

Bjerge K, Nielsen JB, Sepstrup MV, Helsing-Nielsen F, Høye TT. An automated light trap to monitor moths (Lepidoptera) using computer vision-based tracking and deep learning. Sensors (Switzerland) 2021b;doi:10.3390/s21020343.

Caruana R. Multitask Learning. Machine Learning 1997;28(1). doi:10.1023/A:1007379606734.

Dimitrovski I, Kocev D, Loskovska S, Dzeroski S. Hierarchical classification of diatom images using ensembles of predictive clustering trees. Ecological Informatics 2012;7(1). doi:10.1016/j.ecoinf.2011.09.001.

Gao D. Deep Hierarchical Classification for Category Prediction in E-commerce System. In: Proceedings of the 3rd Workshop on e-Commerce and NLP. Association for Computational Linguistics; 2020. p. 64–8. doi:10.18653/v1/2020.ecnlp-1.10.

GBIF. Global Biodiversity Information Facility. 2022. URL: https://www.gbif.org/.

Geissmann Q, Abram PK, Wu D, Haney CH, Carrillo J. Sticky Pi is a high-frequency smart trap that enables the study of insect circadian activity under natural conditions. PLoS Biology 2022;20(7):1–26. doi:10.1371/journal.pbio.3001689.

Golding Y, Ennos R, Sullivan M, Edmunds M. Hoverfly mimicry deceives humans. Journal of Zoology 2005;266(4). doi:10.1017/S0952836905007089.

Gupta DA, Kalhagen ES, Olsen ØL, Goodwin M. Hierarchical Object Detection applied to Fish Species. Nordic Machine Intelligence 2022;2(1):1–15. doi:10.5617/nmi.9452.

Hansen OL, Svenning JC, Olsen K, Dupont S, Garner BH, Iosifidis A, Price BW, Høye TT. Species-level image classification with convolutional neural network enables insect identification from habitus images. Ecology and Evolution 2020;10(2). doi:10.1002/ece3.5921.

He K, Zhang X, Ren S, Sun J. Deep residual learning for image recognition. In: Proceedings of the IEEE Computer Society Conference on Computer Vision and Pattern Recognition. volume 2016-Decem; 2016. doi:10.1109/CVPR.2016.90.

Høye TT, Arje J, Bjerge K, Hansen OLP, Iosifidis A, Leese F, Mann HMR, Meissner K, Melvad C, Raitoharju J. Deep learning and computer vision will transform entomology. Proceedings of the National Academy of Sciences 2021;doi:10.1073/pnas.2002545117.

Kasinathan T, Singaraju D, Uyyala SR. Insect classification and detection in field crops using modern machine learning techniques. Information Processing in Agriculture 2021;8(3). doi:10.1016/j.inpa.2020.09.006.

Kittichai V, Pengsakul T, Chumchuen K, Samung Y, Sriwichai P, Phatthamolrat N, Tongloy T, Jaksukam K, Chuwongin S, Boonsang S. Deep learning approaches for challenging species and gender identification of mosquito vectors. Scientific Reports 2021;11(1). doi:10.1038/s41598-021-84219-4.

van Klink R, August T, Bas Y, Bodesheim P, Bonn A, Fossøy F, Høye TT, Jongejans E, Menz MH, Miraldo A, Roslin T, Roy HE, Ruczynski I, Schigel D, Schaffler L, Sheard JK, Svenningsen C, Tschan GF, Waldchen J, Zizka VM, Astrom J, Bowler DE. Emerging technologies revolutionise insect ecology and monitoring. Trends in Ecology and Evolution 2022;37(10):872–85. doi:10.1016/j.tree.2022.06.001.

Krizhevsky A, Nair V, Hinton G. CIFAR-10 and CIFAR-100 datasets. 2009.

La Grassa R, Gallo I, Landro N. Learn class hierarchy using convolutional neural networks. Applied Intelligence 2021;51(10). doi:10.1007/s10489-020-02103-6.

Lima MCF, Leandro MEDdA, Valero C, Coronel LCP, Bazzo COG. Automatic detection and monitoring of insect pests - A review. Agriculture (Switzerland) 2020;10(5). doi:10.3390/agriculture10050161.

Maurer A, Pontil M, Romera-Paredes B. The benefit of multitask representation learning. Journal of Machine Learning Research 2016;17.

Ong SQ, Hamid SA. Next generation insect taxonomic classification by comparing different deep learning algorithms. PloS one 2022;17(12):e0279094. doi:10.1371/journal.pone.0279094.

Pang G, Shen C, Cao L, Hengel AVD. Deep Learning for Anomaly Detection: A Review. ACM Computing Surveys 2021;54(2). doi:10.1145/3439950.

Park JY, Kim JH. Incremental Class Learning for Hierarchical Classification. IEEE Transactions on Cybernetics 2020;50(1). doi:10.1109/TCYB.2018.2866869.

Preti M, Verheggen F, Angeli S. Insect pest monitoring with camera-equipped traps: strengths and limitations. Journal of Pest Science 2021;94(2):203–17. URL: https://doi.org/10.1007/s10340-020-01309-4. doi:10.1007/s10340-020-01309-4.

Ratnayake MN, Dyer AG, Dorin A. Tracking individual honeybees among wildflower clusters with computer vision-facilitated pollinator monitoring. PLOS ONE 2021;16(2):1–20. URL: https://doi.org/10.1371/journal.pone.0239504. doi:10.1371/journal.pone.0239504.

Redmon J, Farhadi A. YOLOv3: An incremental improvement. 2018. arXiv:1804.02767.

Salakhutdinov R, Tenenbaum JB, Torralba A. Learning with hierarchical-deep models. IEEE Transactions on Pattern Analysis and Machine Intelligence 2013;35(8). doi:10.1109/TPAMI.2012.269.

Sandaruwan PD, Wannige CT. An improved deep learning model for hierarchical classification of protein families. PLoS ONE 2021;16(10 October 2021). doi:10.1371/journal.pone.0258625.

Silla CN, Freitas AA. A survey of hierarchical classification across different application domains. Data Mining and Knowledge Discovery 2011;22(1-2). doi:10.1007/s10618-010-0175-9.

Smith LN. A disciplined approach to neural network hyper-parameters: Part 1 – learning rate, batch size, momentum, and weight decay. arXiv 2018;:1–21URL: http://arxiv.org/abs/1803.09820. arXiv:1803.09820.

Tan M, Le QV. EfficientNet: Rethinking model scaling for convolutional neural networks. In: 36th International Conference on Machine Learning, ICML 2019. volume 2019-June; 2019. .

Taylor S, Jaques N, Nosakhare E, Sano A, Picard R. Personalized Multitask Learning for Predicting Tomorrow’s Mood, Stress, and Health. IEEE Transactions on Affective Computing 2020;11(2). doi:10.1109/TAFFC.2017.2784832.

Terven J, Cordova-Esparza D. A comprehensive review of yolo: From yolov1 and beyond. 2023. arXiv:2304.00501.

Tresson P, Carval D, Tixier P, Puech W. Hierarchical Classification of Very Small Objects: Application to the Detection of Arthropod Species. IEEE Access 2021;9. doi:10.1109/ACCESS.2021.3075293.

Ugenteraan. Deep Hierarchical Classification. 2020. URL: https://github.com/Ugenteraan/Deep_Hierarchical_Classification/; github.

Villa-Perez ME, Alvarez-Carmona M, Loyola-Gonzalez O, Medina-Perez MA, Velazco-Rossell JC, Choo KKR. Semi-supervised anomaly detection algorithms: A comparative summary and future research directions. Knowledge-Based Systems 2021;218. doi:10.1016/j.knosys.2021.106878.

Wu C, Tygert M, LeCun Y. A hierarchical loss and its problems when classifying non-hierarchically. PLoS ONE 2019;14(12):1– 19. doi:10.1371/journal.pone.0226222. arXiv:1709.01062.

Xia D, Chen P, Wang B, Zhang J, Xie C. Insect detection and classification based on an improved convolutional neural network. Sensors (Switzerland) 2018;doi:10.3390/s18124169.

Zhang L, Yang Q, Liu X, Guan H. Rethinking hard-parameter sharing in multi-domain learning. In: 2022 IEEE International Conference on Multimedia and Expo (ICME). Los Alamitos, CA, USA: IEEE Computer Society; 2022. p. 1–6. URL: https://doi.ieeecomputersociety.org/10.1109/ICME52920.2022.9859706. doi:10.1109/ICME52920.2022.9859706.

